# Neuronal CBP-1 is required for enhanced body muscle proteostasis in response to reduced translation downstream of mTOR

**DOI:** 10.1101/2024.03.15.585263

**Authors:** Santina Snow, Dilawar Mir, Zhengxin Ma, Jordan Horrocks, Matthew Cox, Marissa Ruzga, Hussein Sayed, Aric N. Rogers

## Abstract

**Background:** The ability to maintain muscle function decreases with age and loss of proteostatic function. Diet, drugs, and genetic interventions that restrict nutrients or nutrient signaling help preserve long-term muscle function and slow age-related decline. Previously, it was shown that attenuating protein synthesis downstream of the mechanistic target of rapamycin (mTOR) gradually increases expression of heat shock response (HSR) genes in a manner that correlates with increased resilience to protein unfolding stress. Here, we investigate the role of specific tissues in mediating the cytoprotective effects of low translation.

**Methods:** This study uses genetic tools (transgenic *C. elegans*, RNA interference and gene expression analysis) as well as physiological assays (survival and paralysis assays) in order to better understand how specific tissues contribute to adaptive changes involving cellular cross-talk that enhance proteostasis under low translation conditions.

**Results:** We use the *C. elegans* system to show that lowering translation in neurons or the germline increases heat shock gene expression and survival under conditions of heat stress. In addition, we find that low translation in these tissues protects motility in a body muscle-specific model of proteotoxicity that results in paralysis. Low translation in neurons or germline also results in increased expression of certain muscle regulatory and structural genes, reversing reduced expression normally observed with aging in *C. elegans*. Enhanced resilience to protein unfolding stress requires neuronal expression of *cbp-1*.

**Conclusion:** Low translation in either neurons or the germline orchestrate protective adaptation in other tissues, including body muscle.

## 1. Introduction

Loss of protein quality control plays a major role in aging and age-related diseases, including the growing pandemic of protein conformational disorders (1). While frequently considered in the context of neurodegeneration, conformational disorders also contribute to functional decline in skeletal muscle, which exacerbates problems related to muscle wasting that diminish mobility and contribute to frailty (2, 3). Ensuring protein quality control is dependent on the ability to mount a sufficient adaptive response to stress that perturbs proteostasis. Mitigating diminution of proteostasis that occurs with normal aging is an important goal in disease prevention and treatment (4).

Ensuring proper maintenance of proteostasis requires both constitutive and induced activation of cytoplasmic and organelle-specific stress response pathways (5–7). Prominent among these pathways is the cytoplasmic heat shock response (HSR), which governs heat shock factors (HSFs) that drive expression of genes important for restoring proteostasis (8). A review of existing literature indicates that low heat shock gene expression is associated with an increase in cellular senescence, while overexpression is inversely correlated, across a number of tissue types (9). Although it is clear that aging leads to loss of proteostasis (10), the reasons why proper function of this response diminishes with age are not fully understood.

Lowering translation downstream of nutrient sensing improves proteostasis (11–13). The mechanistic target of rapamycin (mTOR) governs cellular nutrient sensing and downregulates the materially and energetically expensive process of mRNA translation when food availability is reduced (14). Individually, restricting mTOR signaling or translation can extend lifespan and improve proteostasis, outcomes that are highly conserved across diverse species (14–17).

Translation controlled downstream of mTOR is regulated by the translation initiation cap-binding complex (CBC). It comprises the eukaryotic translation initiation factor (eIF)4G, the eIF4A RNA helicase, and the 5’ mRNA cap-binding protein eIF4E (18). eIF4G acts as a nexus for translation by bringing in other translation factors and helping recruit the small (40S) ribosomal subunit. Although eIF4E or eIF4G can become limiting for cap-mediated translation, eIF4G is also involved in cap-independent translation and has been shown to be a limiting factor in yeast and worms under nutrient stress or when mTOR signaling is attenuated (19–21). Depletion of eIF4G phenocopies differential translation changes from chemical or genetic inhibition of mTOR in mouse embryonic fibroblasts (22). eIF4G also becomes sequestered in stress granules upon exposure to oxidative or thermal stress in mammalian tissue culture (23, 24). These studies show that changes in expression and availability of eIF4G form part of a conserved adaptive response to enhance survival during periods of nutrient scarcity or other environmental stress. Thus, nutrient status, mTOR, and acute stress regulate expression of the CBC factor eIF4G in diverse animal systems.

In *C. elegans*, changes in proteostasis occur rapidly after entry into adulthood (25) and coincide with chromatin remodeling that decreases robustness of the HSR (26). This aspect of *C. elegans* biology makes it possible to rapidly assess genetic or environmental interventions that modulate age-related changes in proteostasis. Previously, we discovered that genetically attenuating eIF4G reverses proteostatic collapse that occurs early in adulthood in a manner that is partly dependent on *hsf-1* (11), a gene encoding the only HSF in *C. elegans*. Lowering translation coincided with constitutive upregulation of HSR target genes. It also restored robustness of HSR activation to a more youthful level in response to heat stress, a phenomenon we refer to here as HSR priming.

Understanding how to maintain proteostasis for healthy aging is complicated by tissue crosstalk interactions in multicellular systems. This is due to the ability of localized stress responses to influence the activity of proteostasis machinery in distal tissues (27–32), including cell non-autonomous activation of the HSR (29). The means and extent of intercellular signaling are not fully characterized, nor is the connection between translation and proteostasis in distal tissues.

Recently, we showed that antagonistic effects of low translation on growth and longevity were separable by tissue-type and had cell non-autonomous effects on reproduction (33). Lowering translation downstream of mTOR in neurons, germline, or hypodermal tissue increased lifespan and negatively affected reproduction, a trade-off posited by Disposable Soma theory (34) and frequently observed in genetic or environmental interventions that increase lifespan (35). However, lowering translation in body muscle reduced lifespan and increased the rate of development and reproductive output, a complete reversal of the typical trade-offs observed from systemic manipulation, but one in line with trade-offs between decreased motility/energy expenditure and increased reproduction predicted by foraging theory (36–38). Taken together, results led us to wonder whether low translation in specific tissues also controls systemic responses to unfolded protein stress mediated by the HSR.

In the current study, we investigated how nutrient sensing and translation controlled by eIF4G in specific tissues influence proteostasis in *C. elegans*. Results indicate that attenuating translation in neural or germline tissue increases resistance to stress and primes the HSR. Furthermore, we found that low translation in neurons or germline were protective in a body muscle-specific proteotoxicity model and led to increased transcription of muscle structural and regulatory genes, reversing age-related attenuation of expression and improving proteostasis in body muscle. Lastly, we found that neural expression of the transcription factor CBP-1 is required for protective effects of low translation.

## 2. Results

### 2.1 Reduced Translation in Neurons or Germline Primes the HSR and Increases Thermotolerance

Since HSR expression is correlated with somatic protection from perturbations in proteostasis, we sought to first resolve the kinetics of HSR priming upon translation attenuation. For this, we employed RNA interference (RNAi) targeting eIF4G, which is known in *C. elegans* as IFG-1 and encoded by the *ifg-1* gene. This treatment started on the first day of adulthood after all tissues are fully developed and continued for one week. Four HSP genes measured as a proxy for the HSR showed a decrease in the first two days after exposure to *ifg-1* RNAi followed by a steady increase to peak levels by the end of the week (Figure 1A). The delay in HSR induction corresponds with a lack of thermoprotection at two days compared to seven days of low translation observed in a previous study (11). Thus, the time in between the rapid drop in translation and peak in HSP gene expression is considered the period of adaptation to low translation conditions.

**Fig 1.**
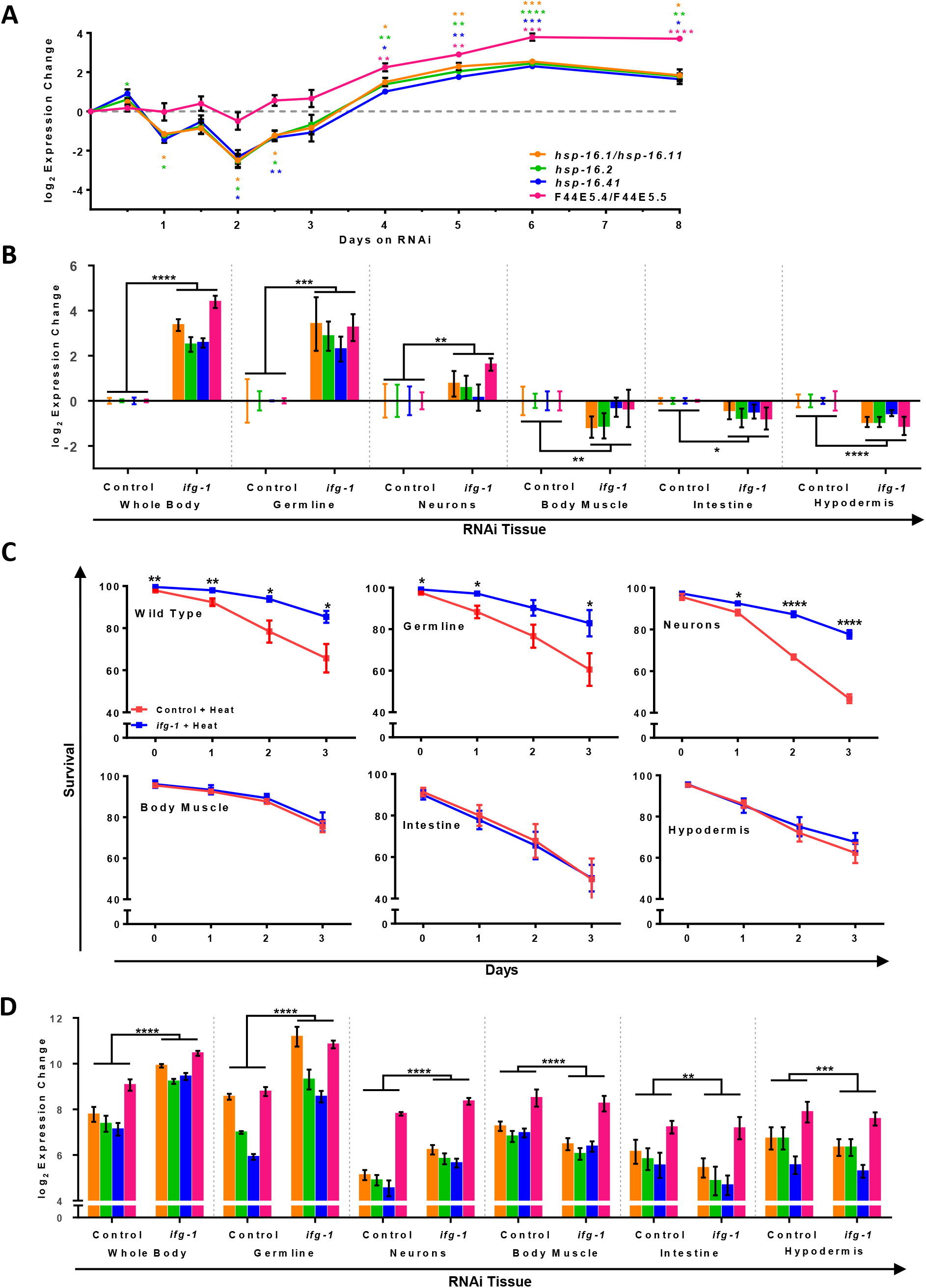
Reduced translation in germline or neurons primes the HSR and increases thermotolerance. (A) Time course of heat shock gene expression in N2 animals under *ifg-1* RNAi normalized to control RNAi at each time point shown, beginning at the onset of adulthood. Unpaired t-tests using Welch’s correction were run on ΔCts for each gene at each time point (Table S3). (B) Heat shock gene expression of tissue-specific strains on *ifg-1* RNAi normalized to control RNAi at day 7 of adulthood. From left to right, panels show N2 wild type animals, MAH23 (germline-specific RNAi strain), TU3335 (neuron-specific RNAi strain), WM118 (body muscle-specific RNAi strain), VP303 (intestine-specific RNAi strain), and NR222 (hypodermis-specific RNAi strain). Two-way ANOVAs were run for each strain comparing ΔCts of all genes (Table S4). (C) Thermotolerance of tissue-specific RNAi strains after 1 week on RNAi. Unpaired t-tests using Welch’s correction were run at each time point for each strain (Table S5). (D) Similar to (B) except that heat shock gene expression was measured 1 hour after exposure to 35°C for 4 hours. Two-way ANOVAs were run for each strain comparing heat shock gene expression (Table S4). Error bars represent means ± SEM. * p<0.05, ** p < 0.01, *** p < 0.001, **** p < 0.0001.

Having established a time-course for HSR priming under *ifg-1* RNAi (hereafter also referred to as a low translation condition), we sought to investigate tissue-specific effects of low *ifg-1* expression. Several *C. elegans* strains were chosen that allow *ifg-1* to be attenuated in select tissues via RNAi as previously characterized (33, 39–42). Tissues selectively targeted by RNAi include the germline, neurons, body muscle, intestine, and hypodermis (for strain details, see methods). Using the same conditions as above, we looked to see whether *ifg-1* RNAi showed signs of HSR priming in these strains. Only low translation in neurons or germline elicited increased HSP expression, whereas other tissues exhibited a very small reduction (Figure 1B, Figure S1). Based on these outcomes, we hypothesized that low translation in neurons or germline tissue would confer protection to thermal stress as observed for whole-body RNAi in a previous study (11).

To determine whether low translation in individual tissues could protect the entire worm from heat-derived unfolded protein stress, wild-type and tissue-specific RNAi strains were treated with *ifg-1* RNAi for one week as in Figure 1B, subjected to thermal stress for four hours, and tracked for survival. Low *ifg-1* expression in N2 wild-type as well as germline or neuronal tissue increased thermotolerance, but did not increase thermotolerance for through other tissues (Figure 1C). Analysis of gene expression showed that only germline- and neuron-specific RNAi strains exhibited more robust HSR gene expression following heat treatment compared to controls (Figure 1D). Findings indicate that either low germline or neuronal translation confers enhanced protection from thermal stress.

Since eIF4G/IFG-1 is downstream of mTORC1, but not mTORC2, we tested both the pathway and tissue-specificity of the response using RNAi targeting Raptor/*daf-15*, a part of the mTORC1 complex and Rictor/*rict-1,* a part of the mTORC2 complex (Figure S2A). *daf-15* RNAi elicited HSR priming in wild-type and germline-specific RNAi animals, but not in the neuronal RNAi strain (Figure S2B). However, enhanced resistance to heat was observed for the neuronal RNAi strain despite the lack of HSR priming (Figure S2B, lower panel). No HSR priming nor protection were conferred by *rict-1* knockdown. Collectively, the data indicates the importance of mTORC1, but not mTORC2 with HSR priming and enhanced thermotolerance.

The transcription factor HSF-1 and the *C. elegans* FOXO transcription factor DAF-16 regulate proteostasis and control stress resistance via the insulin-like signaling pathway (43). To determine whether the molecular alterations and enhanced thermotolerance observed under low translation are dependent on these factors, mutant strains for *daf-16* and *hsf-1* were tested. HSR priming was evident in a *daf-16(mu86)* null mutant in response to *ifg-1* RNAi. Although thermal stress survival tended to be slightly improved, results did not reach statistical significance (Figure S3A). Null mutants for *hsf-1* are not viable and cannot be tested, however, an *hsf-1(sy441)* mutant lacking the carboxy-terminus DNA binding domain was available for testing and failed to induce the same level of HSR priming phenotype, demonstrating the importance of this domain for HSR induction in general (Figure S3B). However, despite greatly reduced HSR priming before heat challenge, following heat treatment, it resulted in a more robust HSR response and improved survival. Interestingly, a previous study showed that overexpression of *hsf-1* lacking the carboxy-terminus (strain AGD794) exhibits enhanced survival (44). Results indicate that the carboxy-terminus of HSF-1 is not necessary for enhanced thermotolerance in general. Collectively, HSR priming and thermotolerance with low translation are partially dependent on the insulin-like signaling pathway.

### 2.2 Reducing Translation in Neurons or the Germline Improves Motility and Restores Youthful Transcription of Muscle Maintenance Genes

The fact that selectively lowering translation in neurons or the germline improve survival from challenge with heat indicates that they are likely to improve maintenance of protein folding in other tissues. Testing this possibility requires a tissue-specific model of protein unfolding stress. For this, we employed motility assays using strain NL5901, which expresses alpha-synuclein fused with YFP in body muscle. This tissue-specific expression is due to the fact that the transgene is controlled by the major heavy myosin chain *unc-54* gene promoter. Aging animals lose the ability to maintain proteostasis, which is highly exacerbated in body muscle in this strain due to expression of alpha-synuclein, resulting in paralysis in worm middle-age (45). Here, we used paralysis to monitor the onset of proteostatic collapse in body muscle.

Prior to looking at tissue-specific effects, we tested the effect of whole body attenuation of *ifg-1* during adulthood, which increased the motile period in NL5901 by an average of 44% (Figure 2A). Unexpectedly, we observed that low translation led to increased alpha-synuclein::YFP fluorescence intensity by the end of the first week of adulthood (Figure 2B,C). Increased protein expression was observed also in a Western blot probed with a monoclonal antibody specific for alpha-synuclein (Figure 2D). Thus, despite increased motility under this condition, lowering systemic translation resulted in increased total protein expression for alpha-synuclein::YFP.

**Fig 2.**
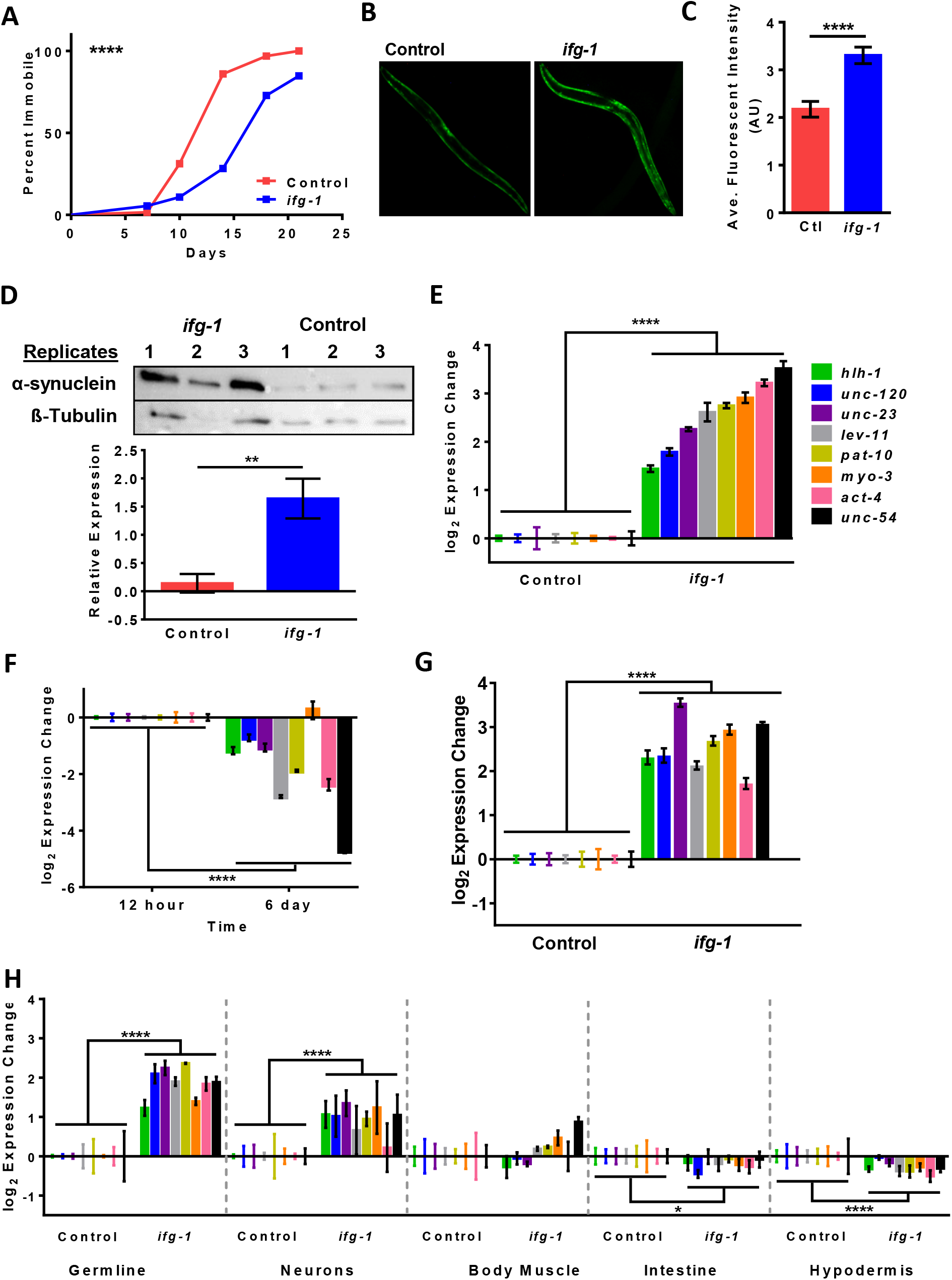
Reducing translation in the germline or neurons improves motility and restores youthful transcription of muscle maintenance genes. (A) Paralysis assay of proteinopathy model expressing alpha-synuclein in body muscle (strain NL5901) on control or *ifg-1* RNAi from the onset of adulthood. Kaplan-Meier survival curves were plotted for paralysis assays and compared using the Mantel-Cox log rank test, p< 0.0001. Additional replicates show similar results (Table S6). (B) Representative images of YFP-tagged alpha-synuclein in NL5901 on control and *ifg-1* RNAi for 7 days. Images were collected from at least 9 animals per replicate across 4 replicates. (C) Quantification and comparison of fluorescence in NL5901 animals from (B) using an unpaired t-test with Welch’s Correction (Table S7). (D) Western blot probing for alpha-synuclein and beta-tubulin protein in NL5901 animals comparing conditions from (B) using an unpaired t-test with Welch’s Correction (Table S8). (E) Expression of muscle structural and regulatory genes in NL5901 animals on control vs *ifg-1* RNAi at d7 of adulthood compared using two-way ANOVA (Table S9). (F) Aging-related changes in muscle-related gene expression of N2 animals on control RNAi using two-way ANOVA, p < 0.0001. (G) Expression of body muscle genes in N2 animals on control or *ifg-1* RNAi from day 1 to day 7 of adulthood compared using two-way ANOVA, p < 0.0001. (H) From left to right, panels show body muscle gene expression on control or *ifg-1* RNAi for the first 7 days of adulthood in MAH23 (germline-specific RNAi strain), TU3335 (neuron-specific RNAi strain), WM118 (body muscle-specific RNAi strain), VP303 (intestine-specific RNAi strain), and NR222 (hypodermis-specific RNAi strain). Two-way ANOVA was used (Table S10). Error bars represent means ± SEM. * p<0.05, ** p < 0.01, *** p < 0.001, **** p < 0.0001.

We wondered whether this result could be explained by increased transcriptional activity of the major heavy myosin chain gene (*unc-54*) promoter used to drive body muscle expression of alpha-synuclein. After seven days of *ifg-1* RNAi started at the onset of adulthood, the transcript level of endogenous *unc-54* was increased by more than 10-fold (Figure 2E). In addition to *unc-54*, we tested expression of several other muscle structural and regulatory genes including muscle actin (*act-4*), minor myosin heavy chain (*myo-3*), troponin C (*pat-10*), tropomyosin (*lev-11*), myogenic transcription factors (*hlh-1, unc-120*), and the *unc-23* gene encoding a negative regulator of proteosomal degradation. Expression of these genes increased in NL5901 subjected to *ifg-1* RNAi. To rule out the possibility of this phenomenon being specific to this strain, we tested wild-type N2 animals with *ifg-1* RNAi. A previous study showed that *unc-54* decreased in the first week of adulthood in *C. elegans* (46). We also observed decreased expression at the end of the first week of adulthood in wild-type animals for *unc-54* and several other muscle structural and regulatory genes (Figure 2F). Transcript expression of these muscle specific genes showed that lowering translation reversed the age-related loss of muscle specific gene expression in wild-type animals (Figure 2G). Tissue-specific RNAi strains demonstrated that increased muscle gene expression resulted from lowering translation in the germline or neurons (Figure 2H). Interestingly, lowering translation selectively in body muscle tissue did not induce body muscle expression significantly (Figure 2H).

Because selectively lowering translation in neurons and germline resulted in HSR priming (Figure 1) and an increase in muscle specific transcripts (Figure 2), we hypothesized that selectively lowering translation in neurons or germline in the alpha-synuclein proteotoxicity model could extend their motile period and delay disease onset. We also wondered what effect lowering translation selectively in body muscle, where proteotoxicity occurs, could improve conditions in this model. Thus, we crossed the corresponding tissue-specific RNAi strains with NL5901. We confirmed that muscle structural and regulatory genes increased when translation was reduced selectively in the germline or neurons (Figure 3A). Interestingly, slightly increased expression of muscle structural and regulatory genes was also observed when translation was selectively lowered in muscle (Figure 3A). This result differs slightly from the results with wild-type N2 animals (Figure 2H), which may be due to the constitutively perturbed muscle proteostasis resulting from expression of alpha-synuclein. The paralysis assay demonstrated that lowering translation in the germline or neuronal tissue resulted in the greatest increase in motility with age, whereas lowering translation in muscle resulted in a small protective effect (Figure 3B). In summary, low translation in the neurons or germline improved body muscle proteostasis.

**Fig 3.**
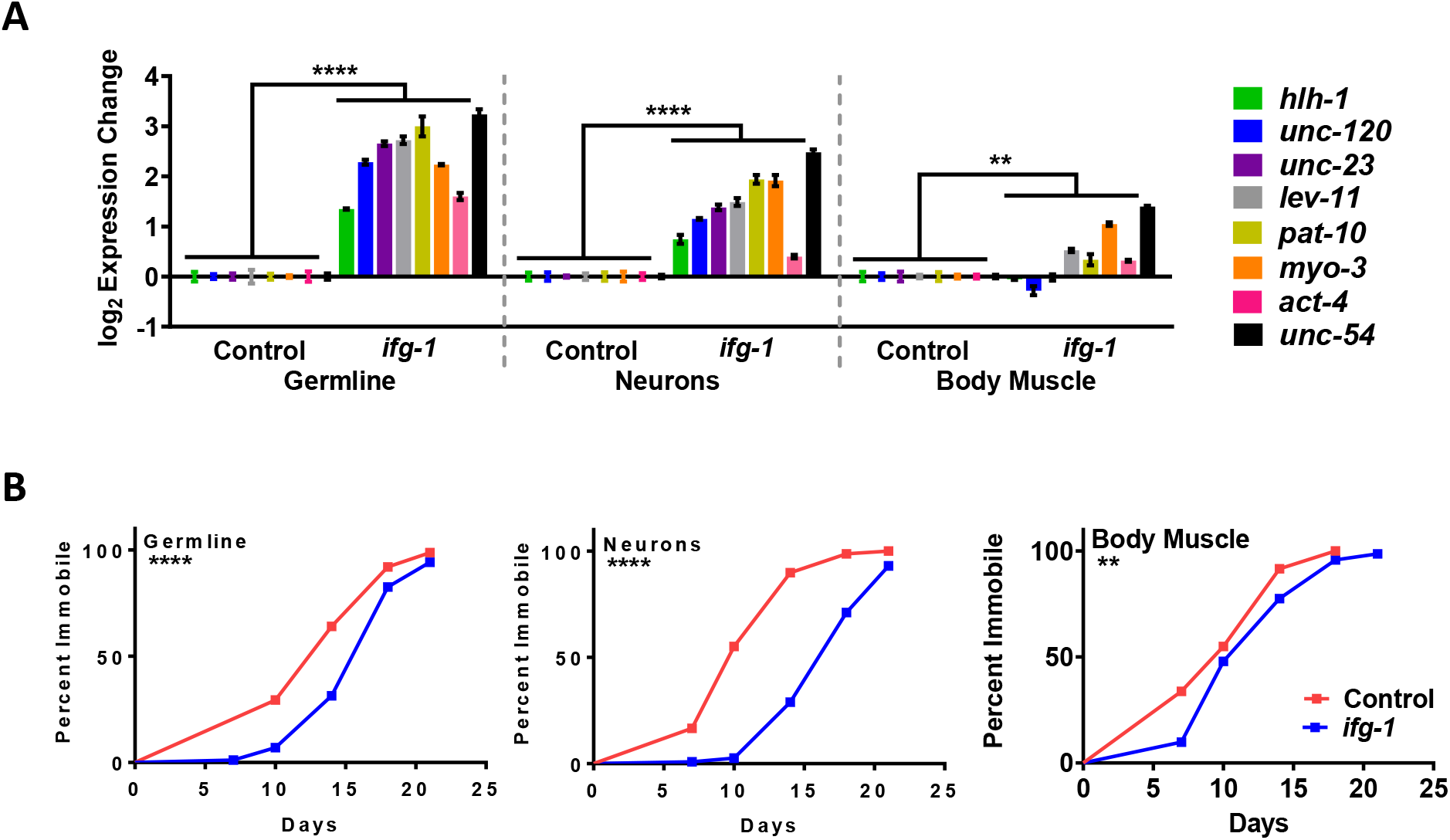
Lowering translation in the germline or neurons upregulates muscle-related gene expression and improves motility in a model of body muscle proteotoxicity. (A) Expression of muscle-related genes after 7 days of *ifg-1* RNAi in the proteinopathy model strain expressing alpha-synuclein in body muscle crossed with tissue-specific RNAi strains (strains ANR149, ANR168, ANR153). Comparisons between animals treated with control or *ifg-1* RNAi were carried out using two-way ANOVAs. (B) Paralysis assays carried out for strains and conditions in (A). Kaplan-Meier survival curves were plotted for paralysis assays and compared using the Mantel-Cox log rank test. From left to right, p< 0.0001, p< 0.0001, and p=0.0089. Replicates were run for each strain with similar results (see Table S6). Error bars represent means ± SEM. * p<0.05, ** p < 0.01, *** p < 0.001, **** p < 0.0001.

### 2.3 Neuronal CBP-1 is Required for Improved Proteostasis Due to Low Translation

We performed a small screen of genes using RNAi to determine whether certain transcription factors, chromatin remodeling factors, or muscle-enriched genes were required for the increase in muscle gene expression observed under low translation conditions. Wild-type N2 animals or neuronal-specific RNAi strain TU3335 were subjected to two days of *ifg-1* RNAi to lower translation before being transferred to another RNAi of interest for an additional 5 days. Translation remained low as no progeny were detected in the days following removal from *ifg-1* RNAi. Because *unc-54* had a robust increase in mRNA transcript abundance under low translation conditions, its expression change was used as a proxy other muscle-specific gene expression changes. Results for a portion of the RNAi tested are shown in Figure S4 and Table S1. From the RNAi screen, the transcriptional regulator gene *cbp-1* was the only gene found to completely suppress the increase in *unc-54* in the neuron-specific RNAi strain, but not in N2 (Figure S4).

Previous studies have shown that *cbp-1* plays an important role in DR-mediated lifespan extension, specifically in neurons (47, 48). Based on this, and its role as a transcriptional regulator, we used the neuron-specific RNAi strain TU3335 and the dual RNAi technique used in the screen to determine whether changes in expression of other body muscle-specific genes were influenced when both *ifg-1* and *cbp-1* expression were reduced in neurons. Results showed that lowering neuronal *cbp-1* expression significantly reduced muscle-specific gene expression changes compared with lowering *ifg-1* by itself (Figure 4A).

**Fig 4.**
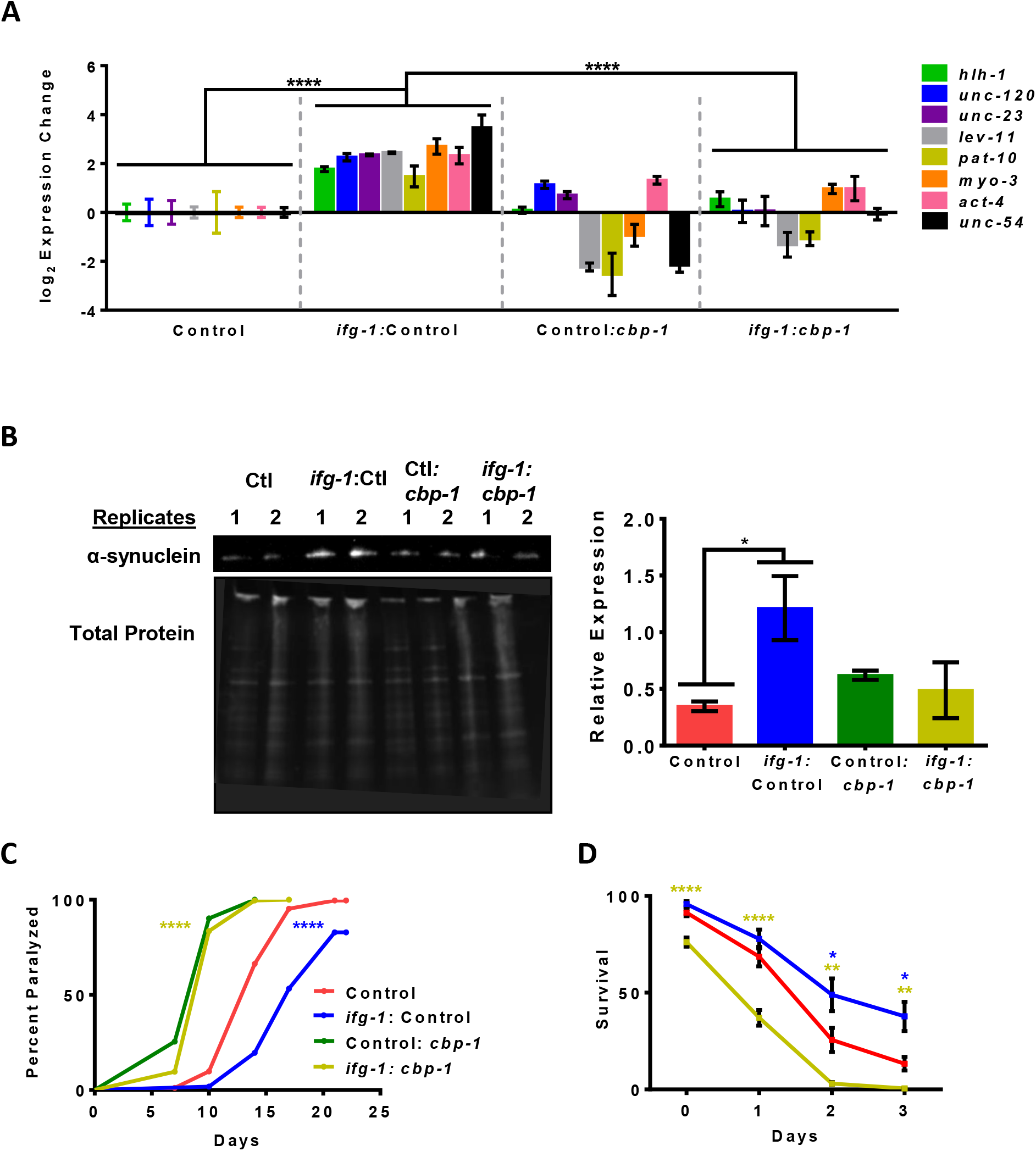
*cbp-1* RNAi in neurons both dampens muscle-related gene expression changes as well as enhanced survival to proteotoxicity under low translation conditions. (A) Expression of genes enriched in muscle in the neuron-specific RNAi strain TU3335 crossed with NL5901 (ANR168). Day 1 adults were placed on control or *ifg-1* RNAi for 2 days before transfer to control or *cbp-1* RNAi for 5 days. Two-way ANOVAs were run comparing ΔCts of the genes shown (Table S11). (B) Western blot of alpha-synuclein (top) and total protein measured from UV shadowing (bottom) for conditions in (A). Quantification and comparison of alpha-synuclein was run using unpaired t-tests with Welch’s Correction (right) (Table S12). (C) Paralysis assay of the strain and conditions from (A). Kaplan-Meier survival curves were plotted for the paralysis assay and compared using the Mantel-Cox log rank test. p < 0.0001 (blue asterisk, ctl vs *ifg1*:ctl) and p < 0.0001 (yellow asterisk, ctl:*cbp-1* vs *ifg-1*:*cbp-1*) (Table S13). (D) Thermotolerance assays (5h at 37°C) using RNAi conditions from (A) in the neuron-specific RNAi strain TU3335. Comparisons at different time points were carried out with unpaired t-tests using Welch’s correction (Table S14). Error bars represent means ± SEM. * p<0.05, ** p < 0.01, *** p < 0.001, **** p < 0.0001.

To determine whether *cbp-1* played a role in the maintenance of muscular proteostasis with low neuronal translation, we crossed the alpha-synuclein proteotoxicity model with neuronal-specific RNAi strain to create ANR168. The combination of *ifg-1* and *cbp-1* RNAi prevented a significant increase alpha-synuclein protein level (Figure 4B). In addition, knocking down expression of neuronal *cbp-1* prevented a robust increase in proteostasis associated with low translation in the muscle paralysis model (Figure 4C). Similarly, *cbp-1* RNAi prevented a robust increase in proteostasis from low translation in the neuronal-specific RNAi strain TU3335 under conditions of heat stress (Figure 4D). Together, results indicate that neuronal CBP-1 is required for the full beneficial effects of low neuronal translation on proteostasis in tissue outside the nervous system, particularly in muscle.

## 3. Discussion

### 3.1 Responses to Low Translation are Partitioned Among Tissues and Capable of Cellular Cross-talk

The basis for loss of proteostatic maintenance with age is centered on inability to properly regulate transcriptional activation of stress response pathways and maintain the protein turnover apparatus (25, 26). Results of this study address the roles of several major tissues in mediating effects of low translation on survival under perturbed proteostasis. Low translation results from downregulation of the mTOR pathway, as occurs when nutrients are scarce, leading to slowed growth but also increased lifespan and resilience to stress. Data support a model in which neurons and the germline are tissues that control physiological responses to low translation, including enhanced somatic maintenance and function of body muscle. Although we do not know how muscle maintenance factors are invoked downstream of translation, we know some of their expression changes are dependent on neuronal expression of *cbp-1*.

CREB-binding factor protein (CBP) acts as a histone acetyltransferase (HAT) to acetylate key transcription factors and histones, and as a recruiter for additional transcriptional machinery (47, 49). Though expressed ubiquitously due to its localization in the nuclei of most somatic cells, in mice hypothalamic expression of CBP is correlated to increases in lifespan (47). At the same time, decreases in its expression are associated with age and diabetes (47). In *C. elegans*, *cbp-1* has been shown to act almost exclusively in GABAergic neurons to double lifespan in DR worms (48). Under DR conditions, its knockdown results in loss of lifespan extension associated with this intervention (47). The current study shows that this factor plays a critical role in adaptive changes downstream of DR and mTOR at the level of translation in neurons.

Based on results from another study, body muscle stands out because it is the only major tissue in which selectively lowering translation reverses, at least in part, the trade-offs usually observed between longevity and development (33). The mTOR pathway is upstream of translation. When low mTOR is driven by nutrient scarcity, it increases mitochondrial respiration in skeletal muscle and mitigates normal respiratory decline observed with age in mice (50). In addition, stem cell availability increases under dietary restriction and low mTOR signaling in injured muscles (51). Thus, it may be that, while translation-inhibiting conditions are detrimental during growth, in adult animals, at least certain aspects of basal muscle maintenance and function are preserved better with age when translation downstream of mTOR is reduced. Our study of the effects of low translation shows an indirect association with enhanced muscle maintenance, requiring low translation in non-muscle tissue. This could help explain the otherwise paradoxical connection between catabolism-inducing dietary restriction or low mTOR signaling conditions and improved muscle maintenance with age.

### 3.2 The Connection between Translational Regulation and Proteostasis

At the heart of improved muscle function and enhanced resistance to unfolded cellular protein resulting from low translation is reversal of age-related diminution of proteostatic maintenance. Many factors are involved in maintaining proteostasis, which involves synthesis, folding, and turnover of cellular protein. Nascent peptide chains emerging from the ribosome are managed by folding chaperones and are subject to ubiquitination, making translation a hub for all these processes (52). The fact that everything from protein synthesis to degradation is regulated at the same location makes translation a critical process for maintaining this balance. We show that limiting protein synthesis controlled by an essential nutrient-responsive translation factor, eIF4G/IFG-1, acts as a lever to restore robustness of stress responsiveness through the HSR. Previously, we showed that at least one other proteostasis mechanism, the ER unfolded protein response, is enhanced when translation is reduced in a manner that depends on the gene encoding the HSR transcription factor, *hsf-1* (53). Experiments with single-celled organisms and mammalian tissue culture showed that intracellular stress signaling pathways can cross-activate one another to maintain cellular proteostasis (30, 54–61). Thus, HSR priming may improve function of other proteostasis mechanisms, a possibility that future experiments will need to address.

## 4. Conclusions

We previously found differential translation on the organismal scale when *ifg-1* was inhibited in a manner consistent with antagonistic modulation between development and somatic maintenance (19). Results in the current study indicate that specific tissues mediate effects of this trade-off to different extents. Future studies analyzing tissue-specific translation changes may help resolve the different effects observed.

## 4. Materials and Methods

### 4.1 Nematode Culture and Strains

*Caenorhabditis elegans* strains were cultured and maintained with standard procedures as described in (62) unless otherwise specified. The N2 Bristol strain was used as the reference wild type. Genotypes of strains acquired from the Caenorhabditis Genetics Center are as follows: MAH23 *rrf-1(pk1417)*, VP303 *rde-1(ne219)*; kbIs7[*nhx-2*p::*rde-1* + *rol-6(su1006)*], NR222 *rde-1(ne219)*; kbIs9[*lin-26p*::nls::GFP, *lin-26p*::*rde-1* + *rol-6(su1006)*], WM118 *rde-1(ne300)*; neIs9[*myo-3*p::HA::*rde-1* + pRF4(*rol-6(su1006)*)], TU3335 *lin-15B(n744)*; uIs57 [*unc-119*p::YFP + *unc-119*p::*sid-1* + *mec-6*p::*mec-6*], PS3551 *hsf-1(syf441),* CF1038 *daf-16(mu86)*; uthIs225 [*sur5p::hsf-1*(CT-Delta)::*unc-54* 3’UTR + *myo-2p*::tdTomato::*unc-54* 3’ UTR], and NL5901 pkIs2386 [*unc-54*p::α-synuclein::YFP + *unc-119*(+)].

Transgenic lines ANR149 *rrf-1(pk1417)*; pkIs2386[*unc-54*p:: α-synuclein::YFP + *unc-119*(+)], ANR153 *rde-1(ne300)*; neIs9[*myo-3*p::HA::*rde-1* + pRF4(*rol-6(su1006)*)]; pkIs2386[*unc-54*p:: α-synuclein::YFP + *unc-119*(+)], and ANR168 *lin-15b(n744);* pkIs2386[*unc-54p::alphasynuclein::YFP + unc-119(+)*]; uIs57[*unc-119p::YFP + unc-119p::sid-1 + mec-6p::mec-6*] were created by crossing NL5901 males with MAH23, WM118 or TU3335 hermaphrodites, respectively. PCR combined with targeted DNA sequencing was performed to validate genotypes.

### 4.2 RNAi Experiments

RNAi knockdown treatments were performed as described in (63). RNAi bacteria strains included empty vector L4440 (Addgene, Cambridge, MA, USA), *ifg-1* (M110.4), *cbp-1* (R10E11.1), *daf-15* (C10C5.6, gifted from Han Lab), and *rict-1* (F29C12.3) (Ahringer library, Source BioScience, Nottingham, UK). RNAi empty vector L4440 is referred to in text as control RNAi. RNAi was carried out from day 0 of adulthood. In Figure 4 and RNAi screens shown in Figure S4, a dual RNAi strategy was used in which day 0 adults were placed on RNAi for *ifg-1* or control for the first two days before being transferred to plates containing RNAi for *cbp-1* or other screen target genes for 5 days prior to analysis. For a full list of RNAis used in screens shown in Figure S4, refer to Table S1. For the RNAi screen, animals were synchronized via bleaching gravid animals to obtain similarly staged embryos.

### 4.3 Thermotolerance

Approximately 120 synchronized animals were maintained for each condition at 20°C until exposure to heat stress (35°C for 4 hours or 37°C for 5 hours). Animals were allowed to recover for 1 hour before being scored for survival. Thereafter, survival was scored at daily intervals.

### 4.4 RNA processing and qPCR

RNA was isolated using Trizol Reagent following the manufacturer’s instructions. RNA samples were then processed with either Sureprep RNA Cleanup and Concentration Kit (Fisher BioReagents, Fair Lawn, NJ, USA) or RNA Clean and Concentrator (Zymo Research, Irvine, CA, USA). 200ng RNA was reverse transcribed using QuantiTect Reverse Transcription Kit (Qiagen, Valencia, CA, USA). qRT-PCR was performed in technical duplicate or triplicate using SYBR FAST qPCR Master Mix (Kapa Biosystems, Cape Town, South Africa) on a LightCycler 480 (Roche Applied Science, Indianapolis, IN). Target gene mRNA expression was normalized to the housekeeping gene *cdc-42* expression. Relative expression was determined by normalizing to control samples. Primer sequences are provided in Table S2.

### 4.5 Paralysis Analysis

Synchronized NL5901, ANR149, ANR153, and ANR168 strains were maintained on control or *ifg-1* RNAi plates starting at adulthood and transferred fresh RNAi plates daily. On days of paralysis measurement, adults were transferred to a new RNAi plate with their bodies aligned in a straight line. After 10 minutes, worms that had not moved from their original location were gently tapped with a sterile platinum wire 2-3 times on the head. Worms able to move only their head (from the pharynx bulb to the tip of the head) were scored as paralyzed. Worms able to move between the pharynx and tail were considered not paralyzed. Worms unable to respond to touch were scored as dead. Paralyzed or dead worms were removed from the plate on the day of paralysis measurement.

### 4.6 Western Blotting

Synchronized NL5901 and ANR168 strains were maintained as they were in paralysis assays until day 7 when total protein extraction occurred. To determine the levels of alpha-synuclein proteins, western blotting was performed in triplicate. Total protein extraction and preparation was performed as previously described in (11). Alpha-synuclein was detected using an anti-alpha-synuclein mouse monoclonal antibody (1:500 dilution) (Santa Cruz biotechnology, Santa Cruz, CA, USA). Beta-tubulin was detected using an anti-beta-tubulin mouse monoclonal antibody (E7, 1:500 dilution) (DSHB, Iowa City, IA). Peroxidase-conjugated goat anti-mouse IgG secondary antibody (1:5000 dilution) was from Pierce (Rockford, IL. USA). The density of the bands was determined using ImageJ software and normalized according to beta-tubulin or total protein.

### 4.7 Imaging

Synchronized NL5901 animals were maintained were maintained on control or *ifg-1* RNAi plates starting at adulthood and transferred fresh RNAi plates daily. Visualization and quantification of alpha-synuclein::YFP expression was conducted after 7 days of control or *ifg-1* RNAi exposure. Individual worms were mounted on a 2% agarose pad and immobilized in a drop of 25mM levamisol. Worms were imaged on a Leica M 165 FC microscope with the YFP filter (excitation 510/20 nm, emission 560/40nm). The intensity of YFP fluorescence was measured in ImageJ by closely tracing around individual worms. In total, approximately 30-40 worms were used per condition.

### 4.8 Statistical Analysis

All statistics were performed using GraphPad Prism 6 software. Kaplan-Meier survival curves were plotted for paralysis assays and compared using the Mantel-Cox log rank test. Western blots, survival assays, and fluorescent expression analyses were compared using unpaired two-tailed t-tests with Welch’s correction. Data from quantitative PCR were assessed by performing two-way ANOVA or unpaired two-tailed t-tests with Welch’s correction.

## Supporting information

Raw Western Blot Images

## Acknowledgements

This work was supported by grants from the National Institutes of Health (R01AG062575) and by the Morris Scientific Discovery Fund. Research reported in this publication was also supported by an Institutional Development Award (IDeA) from the National Institute of General Medical Sciences of the National Institutes of Health under grant numbers P20GM0103423 and P20GM104318. Some strains were provided by the CGC, which is funded by NIH Office of Research Infrastructure Programs (P40 OD010440). The authors declare no conflicts of interest.

## Supplemental Figure legends

**Fig S1.**
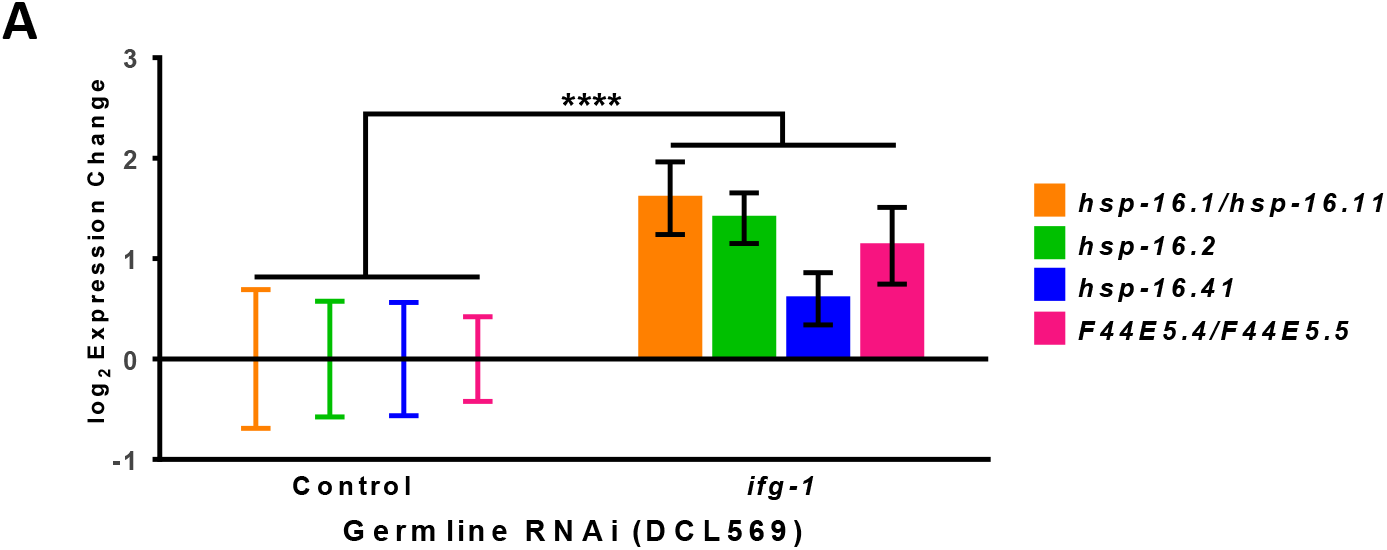
HSR priming using another germline-specific RNAi strain. (A) Heat shock gene expression of the germline-specific RNAi strain (DCL569) after placing adults on control or *ifg-1* RNAi for 7 days. A two-way ANOVA was run comparing ΔCts of all genes. Error bars represent means ± SEM. * p<0.05, ** p < 0.01, *** p < 0.001, **** p < 0.0001.

**Fig S2.**
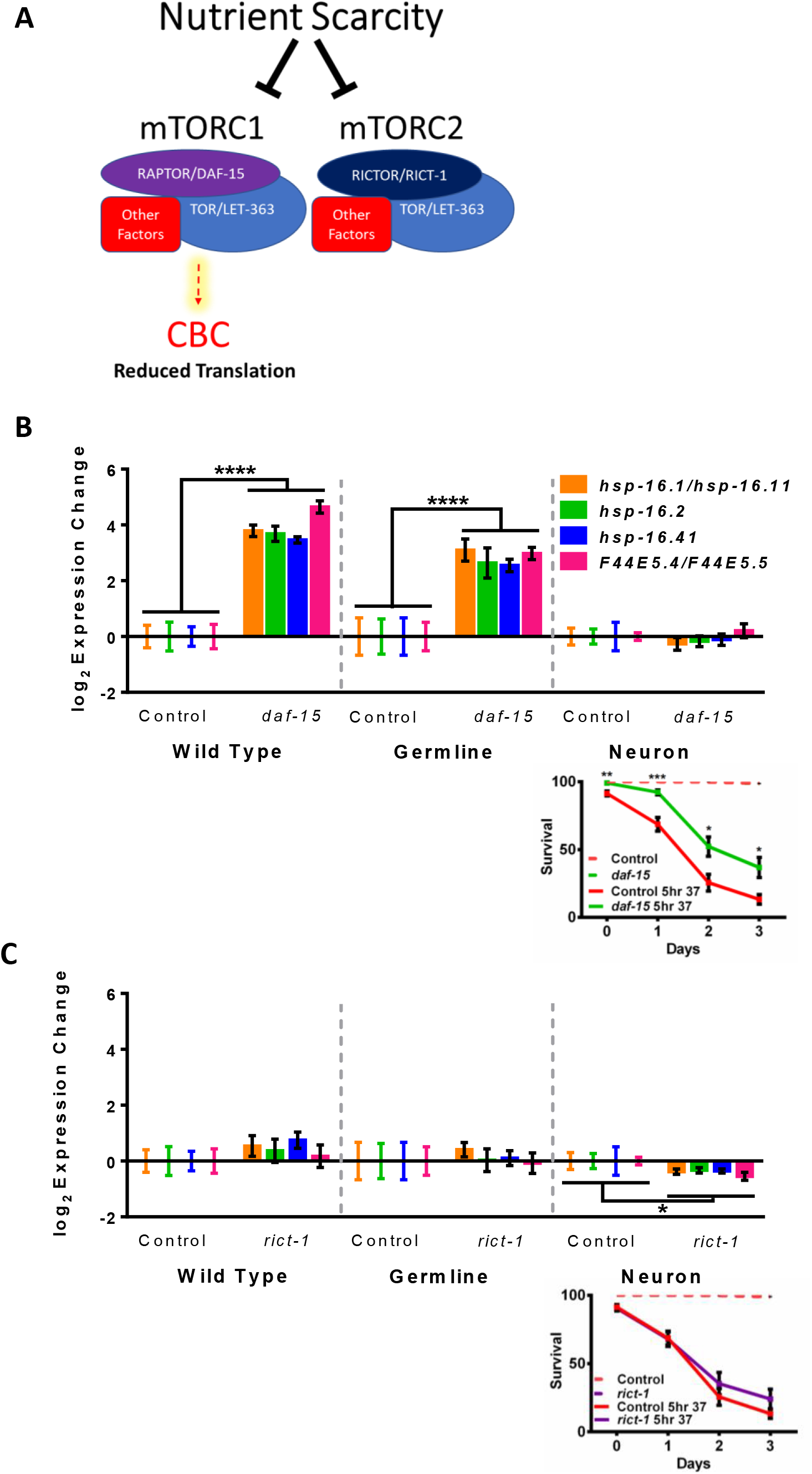
HSR priming through the mTOR pathway. (A) Schematic showing the mTORC pathways inhibited by nutrient scarcity. Only mTORC1 regulates translation, particularly through the Cap-Binding Complex (CBC). (B) Upper panel shows HSR gene expression upon 7 days of exposure to control or *daf-15*/Raptor RNAi of wild-type N2, germline-specific RNAi strain MAH23, and neuron-specific RNAi strain TU3335. Two-way ANOVAs were run for each strain comparing ΔCts of all genes (Table S15). Lower panels show survival subsequent to heat exposure (5h at 37°C). Unpaired t-tests using Welch’s correction were run at each time point in the survival curves (Table S16). (C) Same as in (B) except for *rict-1*/Rictor RNAi as the test. Error bars represent means ± SEM. * p<0.05, ** p < 0.01, *** p < 0.001, **** p < 0.0001.

**Fig S3.**
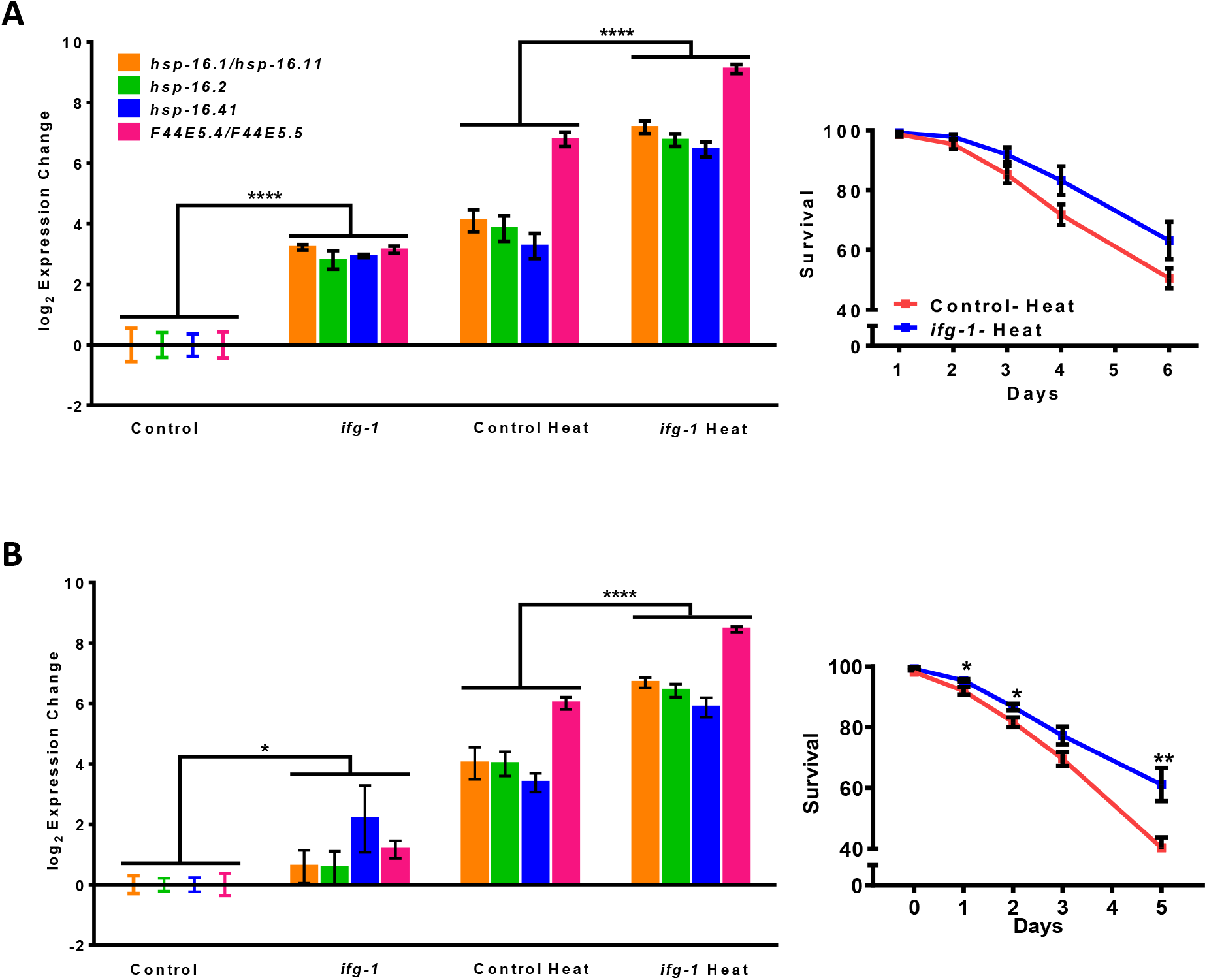
HSR response and thermotolerance in mutants of *daf-16* and *hsf-1*. (A) Left panel shows heat shock gene expression of *daf-16 (mu86)* animals on control and *ifg-1* RNAi for 7 days with and without thermal stress (4h at 35°C). Two-way ANOVAs were run for each strain comparing ΔCts of all genes. Right panel shows survival of *daf-16 (mu86)* animals subjected to heat stress (4h at 35°C). Unpaired t-tests using Welch’s correction were run at each time point (Table S17). (B) Same as in (A), but for *hsf-1 (sy441)* (Table S17). Error bars represent means ± SEM. * p<0.05, ** p < 0.01, *** p < 0.001, **** p < 0.0001.

**Fig S4.**
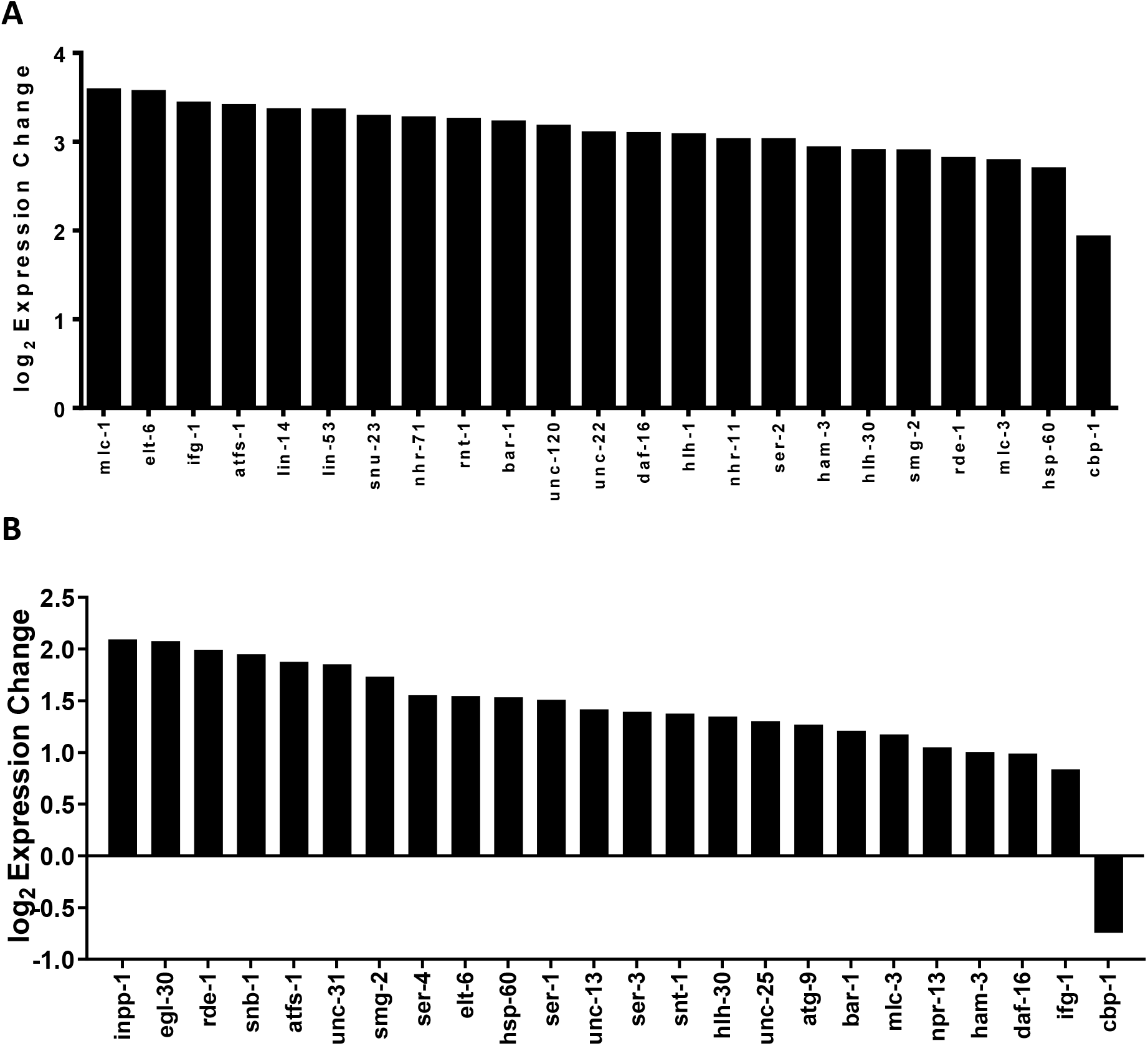
RNAi screen for genes that subdue muscle gene expression changes under low translation conditions. (A) *unc-54* expression of wild-type N2 animals who spent 2 days on *ifg-1* RNAi before being transferred to the RNAis shown for an additional 5 days. (B) Same as in (A) except that the screen was carried out using neuron-specific RNAi strain TU3335. See Supplementary Table S1 for full RNAi gene list used in both strains.

