## Supplementary material for "Neuronal CBP-1 is required for enhanced body muscle proteostasis in response to reduced translation downstream of mTOR": Raw Western Blot Images

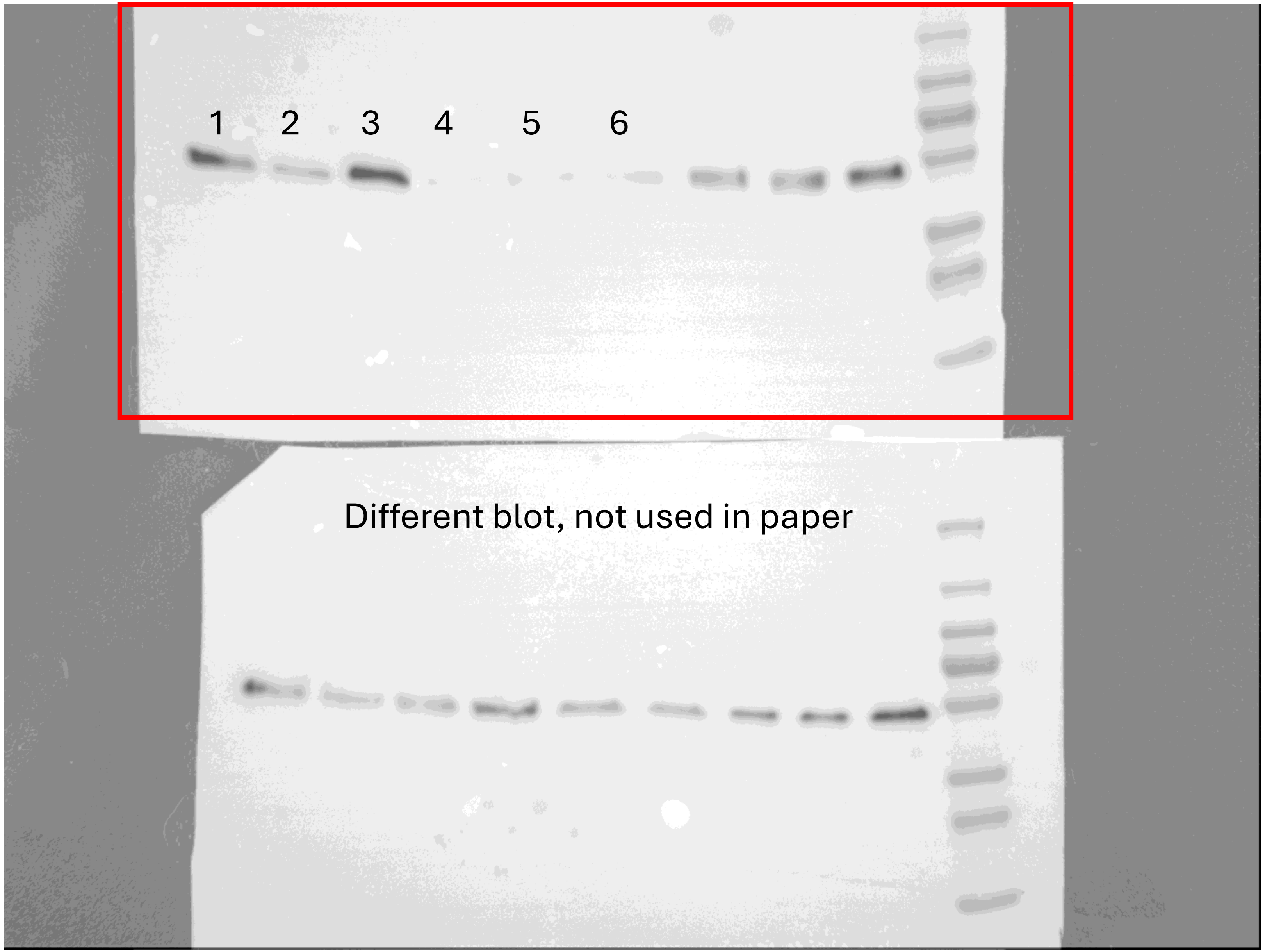


FIGURE 2D
Alpha-synuclein probe blot results

1- ifg1 1

2-ifg-1 2

3- ifg-1 3

4- ctl 1

5- ctl 2

6- ctl 3

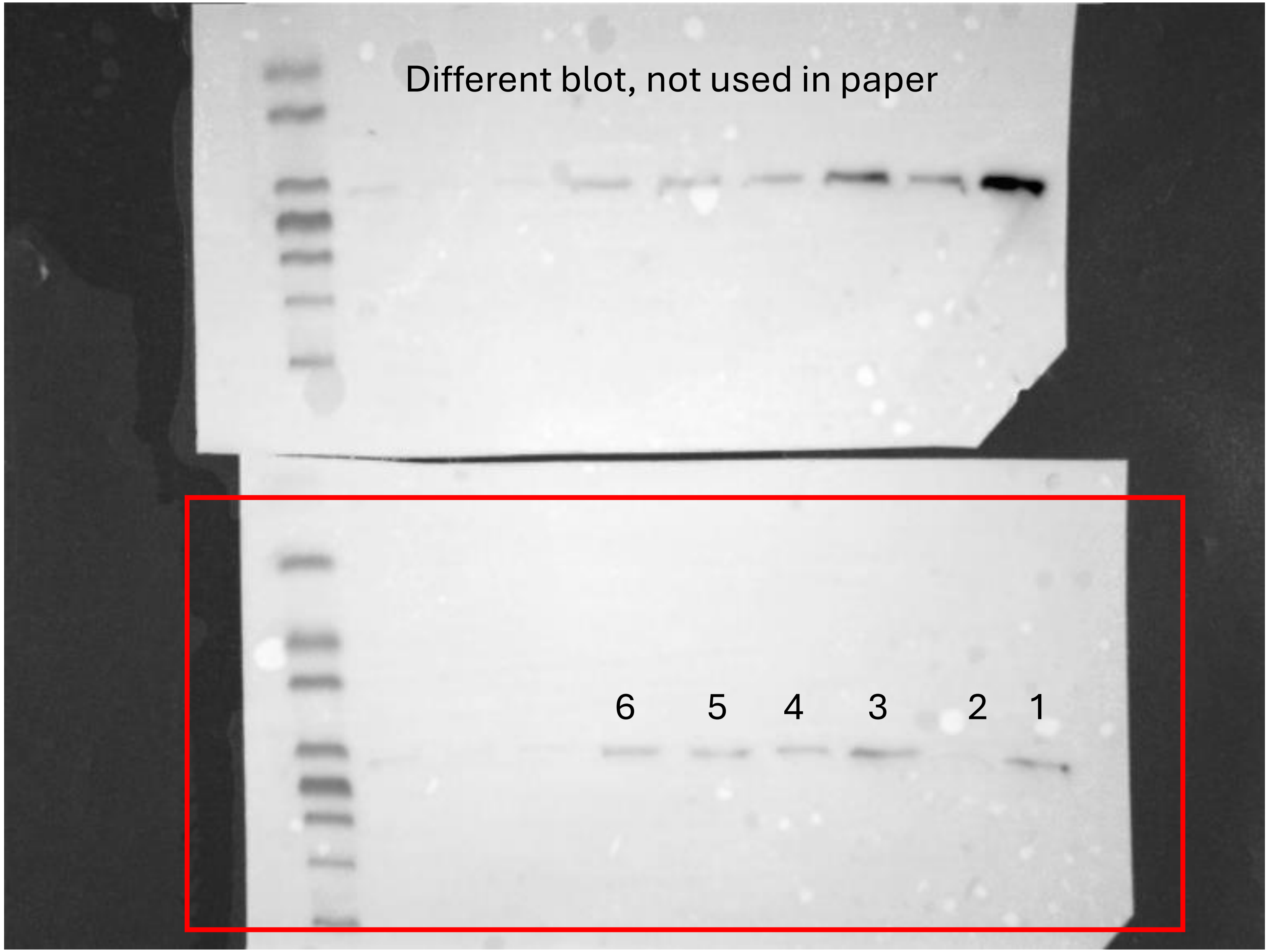


FIGURE 2D

Beta-tubulin probe blot results

1- ifg1 1

2-ifg-1 2

3- ifg-1 3

4- ctl 1

5- ctl 2

6- ctl 3

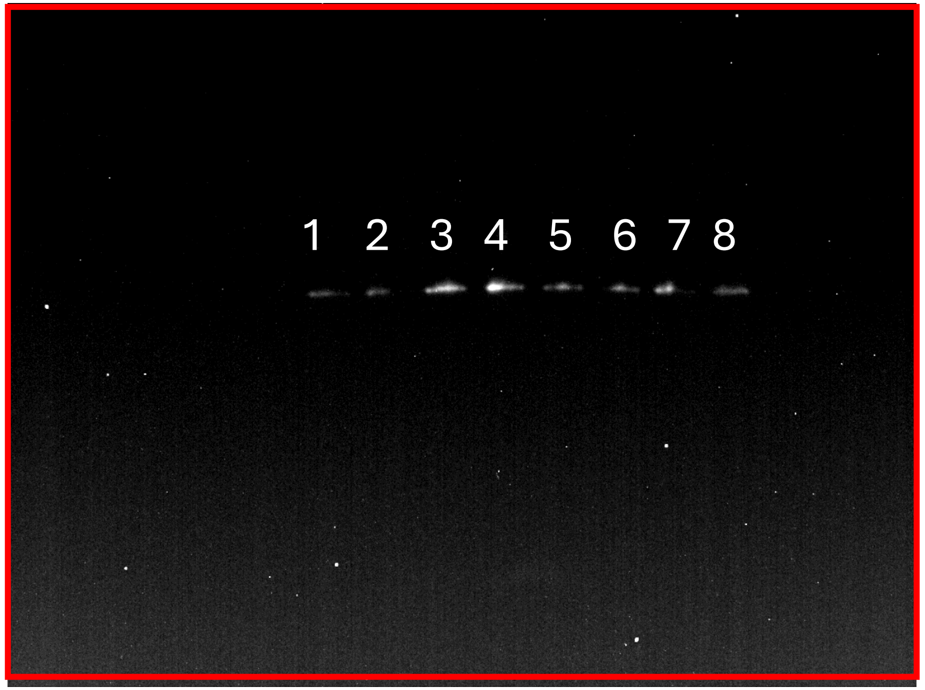


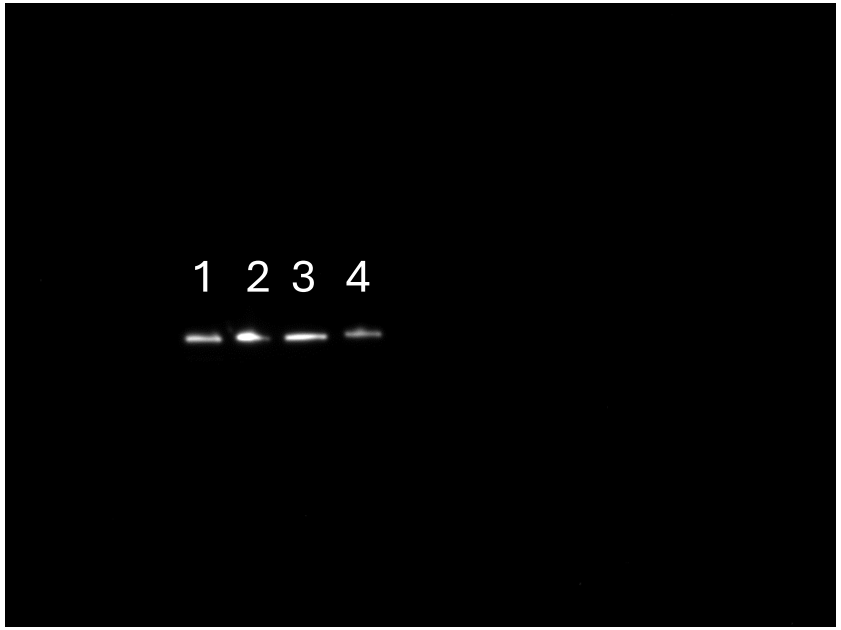


FIGURE 4B Alpha-synuclein probe blot results

Shown in paper:

1-ctl 1

2-ctl 2

3- ifg-1:ctl 1

4- ifg-1:ctl 2

5-ctl:cbp-1 1

6-ctl:cbp-1 2

7-ifg-1:cbp-1 1

8-ifg-1:cbp-1 2

Not shown in paper, but used to quantify 4B:

1-ctl 3

2-ifg-1:ctl 3

3-ctl:cbp-1 3

4-ifg-1:cbp-1 3

FIGURE 4B Total protein gel results

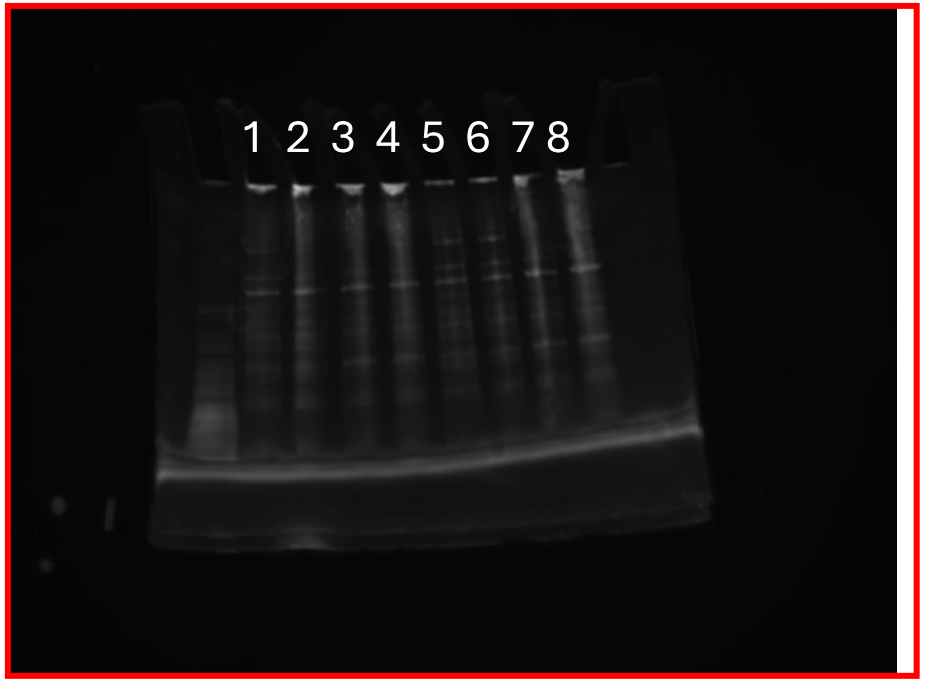


Shown in paper:

1-ctl 1

2-ctl 2

3- ifg-1:ctl 1

4- ifg-1:ctl 2

5-ctl:cbp-1 1

6-ctl:cbp-1 2

7-ifg-1:cbp-1 1

8-ifg-1:cbp-1 2

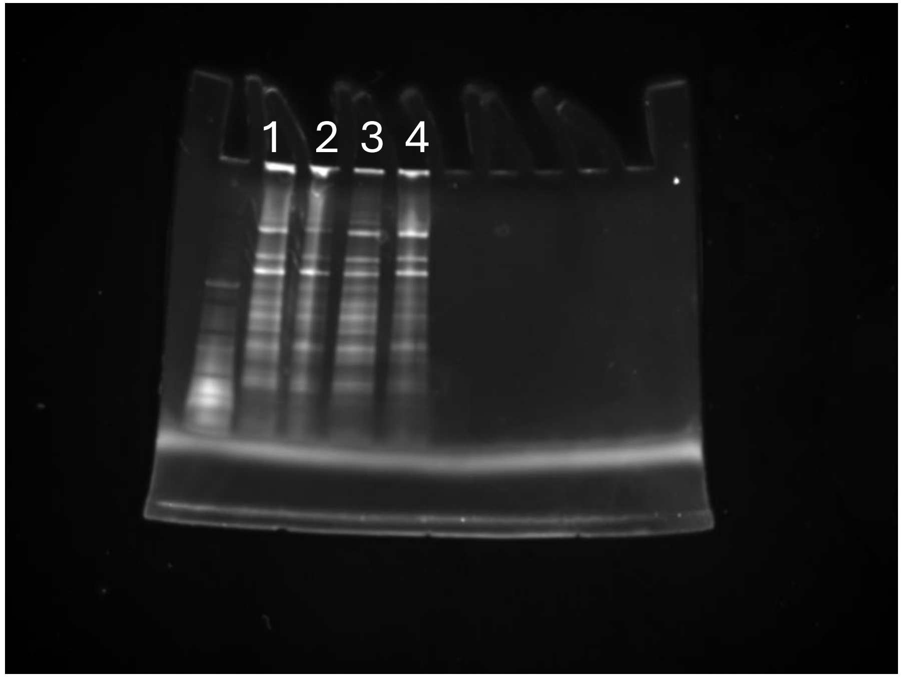


Not shown in paper, but used to quantify 4B:

1-ctl 3

2-ifg-1:ctl 3

3-ctl:cbp-1 3

4-ifg-1:cbp-1 3
